# Identifying disease genes using machine learning and gene functional similarities, assessed through Gene Ontology

**DOI:** 10.1101/472217

**Authors:** Muhammad Asif, Hugo F. M. C. M. Martiniano, Astrid M. Vicente, Francisco M. Couto

## Abstract

Identifying disease genes from a vast amount of genetic data is one of the most challenging tasks in the post-genomic era. Also, complex diseases present highly heterogeneous genotype, which difficult biological marker identification. Machine learning methods are widely used to identify these markers, but their performance is highly dependent upon the size and quality of available data.

In this study, we demonstrated that machine learning classifiers trained on gene functional similarities, using Gene Ontology (GO), can improve the identification of genes involved in complex diseases. For this purpose, we developed a supervised machine learning methodology to predict complex disease genes. The proposed pipeline was assessed using Autism Spectrum Disorder (ASD) candidate genes. A quantitative measure of gene functional similarities was obtained by employing different semantic similarity measures. To infer the hidden functional similarities between ASD genes, various types of machine learning classifiers were built on quantitative semantic similarity matrices of ASD and non-ASD genes. The classifiers trained and tested on ASD and non-ASD gene functional similarities outperformed previously reported ASD classifiers. For example, a Random Forest (RF) classifier achieved an AUC of 0. 80 for predicting new ASD genes, which was higher than the reported classifier (0.73). Additionally, this classifier was able to predict 73 novel ASD candidate genes that were were enriched for core ASD phenotypes, such as *autism* and *obsessive-compulsive behavior*. In addition, predicted genes were also enriched for ASD co-occurring conditions, including Attention Deficit Hyperactivity Disorder (ADHD).

We also developed a KNIME workflow with the proposed methodology which allows users to configure and execute it without requiring machine learning and programming skills. Machine learning is an effective and reliable technique to decipher ASD mechanism by identifying novel disease genes, but this study further demonstrated that their performance can be improved by incorporating a quantitative measure of gene functional similarities. Source code and the workflow of the proposed methodology are available at https://github.com/Muh-Asif/ASD-genes-prediction.

## Introduction

Complex diseases with a strong genetic influence, such as Autism Spectrum Disorder (ASD), often have multiple etiologies with the involvement of possibly hundreds of different genes. Recent advances in genetic technologies have led to more efficient acquisition of genetic data. For example, the many large-scale genetic studies for ASD have identified hundreds of candidate disease (1,2). However, given the vast amount of data generated by these large-scale studies, identifying true disease genes has become a daunting task. For example, a genetic etiology is identified for only 20 to 25% of ASD cases (3), thus demanding more efforts to unravel ASD causing genes.

Random walk with restart (RWR) algorithm has been widely used for disease gene prediction (4,5). Recently machine learning methods have become a powerful computational technique in analyzing genomic datasets (6). Supervised machine learning methods trace hidden relationships among disease-causing genes in existing datasets, such as gene co-expression profiles, functional similarities, or protein-protein interactions networks; and then uses this information to discriminate disease genes from non-disease genes (7–9). Machine learning methods are helping biologists to define genomic features that could have a role in disease mechanism. Krishnan et al. (10) reported a weighted Support Vector Machine (SVM) classifier to predict the probability of association of each brain gene with ASD. The weighted SVM was trained on a specialized gene network that integrates gene expression, protein-protein interactions, and regulatory-sequence information of brain genes. However, classifier evaluation was limited to a subset of genes that were high confidence ASD genes. Such held out restricted evaluation of classifiers may overestimate classifier performance. Moreover, there are limitations for protein-protein interactions and co-expression networks. For example, it is difficult to map weak interactions on interactions networks, thus, limiting the scope of disease gene prediction. To overcome the shortcomings of network-based annotations, studies have used Gene Ontology (GO) (http://www.geneontology.org/) (11), which is a highly efficient resource for predicting disease-causing genes (12).

Functionally similar genes tend to contribute to similar phenotype. For instance, etiologically relevant genes disrupted by genetic variants in ASD patients tend to aggregate in specific biological processes (13), indicating that disease genes may belong to the same hierarchical path of GO and will have higher functional similarities. Therefore, functional similarity of genes, measured using GO, can also be used to predict potential ASD candidate genes.

In this study, we hypothesized that machine learning methods trained and tested only on GO-based gene functional similarities can be used to predict disease-associated genes with improved performance over state of the art methods (10). We validated the performance of classifiers using the same assessment approach as done by Krishnan et al., namely held-out restricted cross-validation. In addition, classifiers were also validated by stratified five-fold cross-validation. Different machine learning methods were applied to a functional similarity matrix of known ASD and non-mental genes. Classifiers trained and tested only on gene functional similarities performed better than the state of the art method. This shows that GO functional similarity is a relevant resource to predict ASD genes. Thus, machine learning methods trained and tested on gene semantic similarities can effectively narrow down the genetic complexity of ASD. Additionally, a classifier built on high confidence ASD genes obtained improved performance, and hence could be used for ASD gene prediction. We also presented a fully automated and customizable KNIME workflow of the proposed methodology, which can be used to identify any other type of disease genes.

## Methods

### Overview of the proposed methodology

In this study, we developed a machine learning based methodology for the identification of disease genes. Fig 1 shows the generic pipeline of the proposed methodology. For any given set of disease-relevant and non-relevant genes, a functional similarity matrix is constructed (step 1). Functional similarities between genes can be determined using gene expression profiles, protein-protein interactions networks, or GO. Different machine learning algorithms are trained and tested on functional similarity matrices (step 2 and 3). To avoid biased estimates, a reasonably balanced ratio of disease and non-disease genes is required. An easy to use and automated workflow of the proposed methodology, using the KNIME machine learning framework, is also provided for the users with minimal programming or machine learning skills.

**Fig1.**
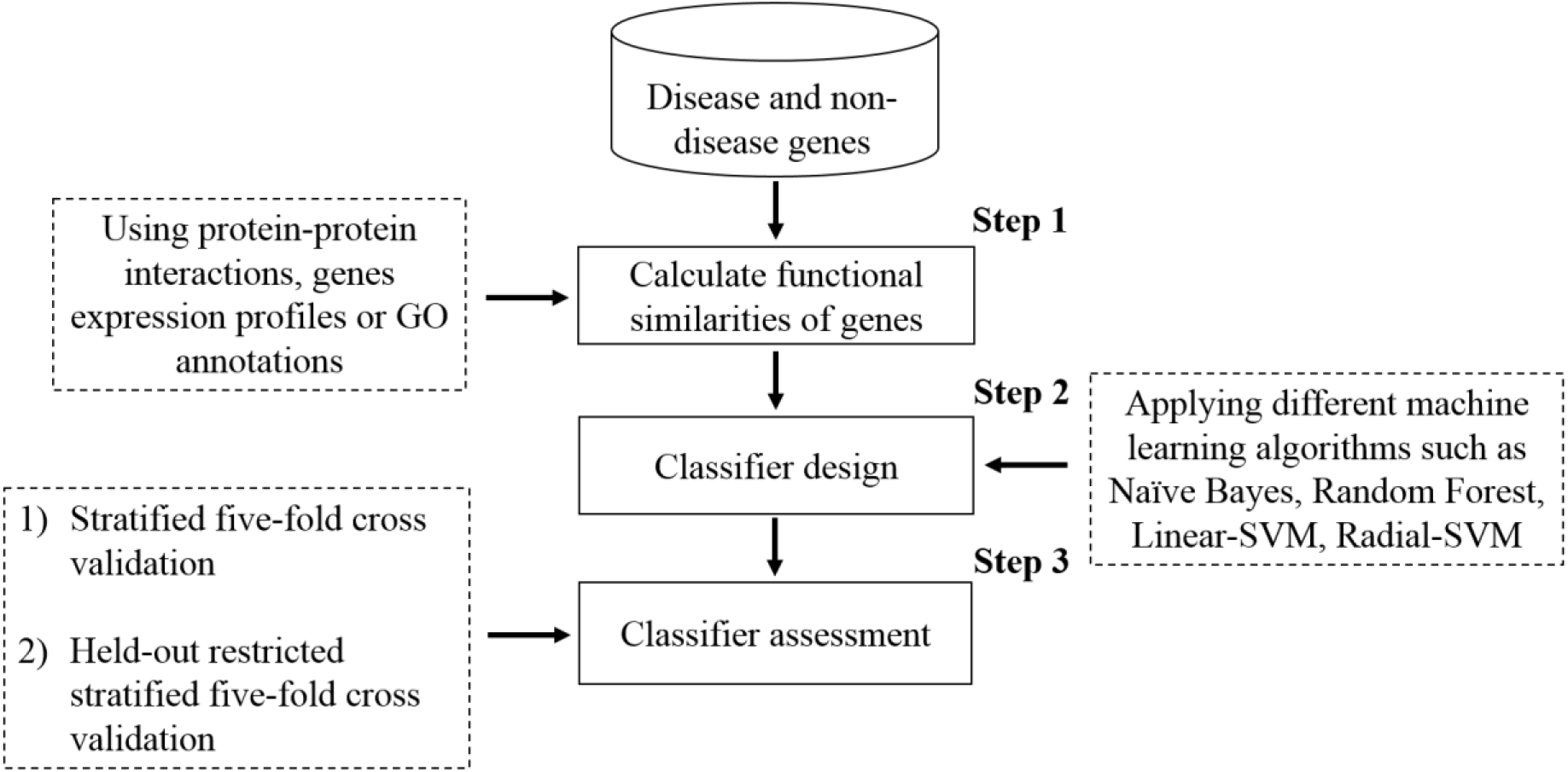
Main components of the proposed methodology to predict disease genes. Functional similarities are computed for a given set of genes. Different machine learning methods are applied to functional similarity matrices to define rules that discriminate disease genes from non-disease genes. Two evaluation approaches, namely stratified and held-out restricted stratified five-fold cross-validation are used (more detail is depicted in Fig 3).

### Machine learning methods

For this study, we used different machine learning classification methods, such as Naive Bayes (NB) (14), linear and radial SVM (15), and a decision tree based method, Random Forest (RF) (16). Supervised machine learning methods have been widely used to predict disease genes. A comparison of classification based methods can be found in Le et al. (17).

RF, linear and radial SVM, and NB were trained on functional similarities of disease and non-disease genes. RF is an ensemble learning method and builds a forest of decision trees. Each tree in forest is a separate weak model that is trained with a randomly selected subset of data and each node of the tree is split by random selection of features from training data. This randomness permits RF to create many uncorrelated weak models, which are then combined to generate a robust and stable model. In the case of classification, the final class attributions from RF are the result of consensus voting from all weak models. RF was applied using the following parameters: number of trees (ntree) = 500; number of variables randomly selected at each split (mtry) equal to the square root of the total number of features (Breiman et al.’s recommended value), random sampling with replacement; and equal class weights assumed for both positive and negative classes. We used the RF implementation from the randomForest R package (version 4.6-12) (18).

The SVM is a heuristic optimization method that tends to discover coordinates of observations, called support vectors, to determine a decision hyperplane that best segregates both classes (positive and negative) with maximum margins in an n-dimensional space, where n is the number of features used to model. SVM can be used as a linear or non-linear method by defining kernel function. In non-linear SVM the dot products of linear SVM are replaced with kernel function that allows the algorithm to map data in higher dimensional feature space. In this study, we used linear SVM and radial kernel-based SVM to predict disease genes. Linear and radial SVM was applied using e1071 (version 1.6.8) R package (19). Equal class weights were defined for both positive and negative classes. 1 and 0.001 values were used for cost and epsilon parameters respectively. For radial-SVM gamma was set to 0.02.

NB is a simple probabilistic classifier based on the Bayes’ theorem with the assumption of strong (naive) independence between every pair of predictors. This simplest Bayesian classifier can be trained efficiently and performs well in practice, even when the assumption of features independence is not valid. It was trained as a conditional probability model, where conditional probability distribution of each observation is estimated using Bayes’ theorem and then uses it for the prediction of class labels for test set instances. NB was applied using e1071 R package. NB was used without Laplace smoothing.

### Evaluation of the classifiers

All classifiers were evaluated using two different assessment approaches, namely stratified five-fold cross-validation and held-out restricted stratified five-fold cross-validation, adapted from the Krishnan et al. study. Stratified five-fold cross-validation consists of following steps:

1. Split the dataset into five equal folds with class probabilities similar to the original dataset
2. Train classifier on four randomly selected folds (training set)
3. Test the trained classifier using the remaining fold (test set).
4. Repeat the process five times and each time a different fold is used as a test set (Fig 3).

**Fig3.**
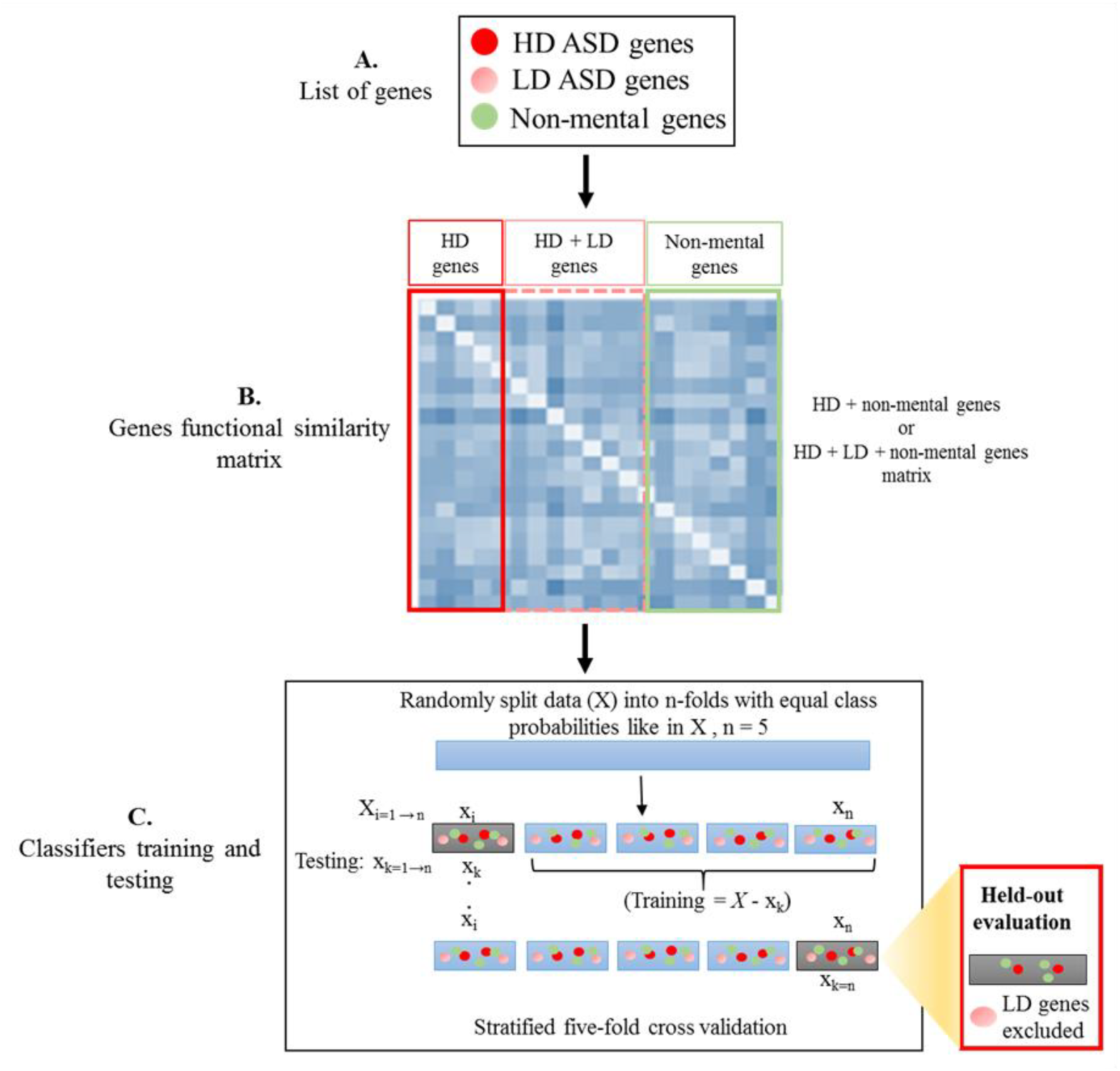
Graphical representation of the methodology to predict ASD genes. **A.** ASD genes with different level of evidence and non-mental genes were used to implement the proposed methodology. **B.** Three different semantic similarity measures were used to calculate functional similarities for HD + non-mental genes and for HD + LD + non-mental genes. **C.** Four different machine learning methods were used to analyze the computed gene functional similarities. Machine learning classifiers were tested using stratified and held-out restricted stratified five-fold cross-validation. Like the Krishnan et al. method in the held-out restricted validation, only HD + non-mental genes were chosen for testing the classifier. In stratified five-fold cross-validation, classifiers were evaluated using all genes in the test set.

In held-out restricted stratified five-fold cross-validation, the classifier is tested using a subset of test set (step 3), named as held-out (Fig 3).

Similar to Krishnan et al.’s method, the Area Under receiver-operator Curve (AUC) evaluation metric was used to estimate and compare the performance of all generated classifiers. To avoid bias from self-similarity scores, during cross-validation, the semantic similarity squared matrix was split into five folds. So, genes selected to be as instances in the test set were removed as features from the test and training set. This avoided the usage of self-similarity values in the test set. For example, for a given set of five genes: g1…g5, the semantic similarity matrix is of 5x5 dimensions. If g1…g3 are used as instances in the training set then g4 and g5 are in the test set so they cannot be used as features in the training set. The training set matrix is therefore reduced to 3x3 dimensions with the three genes: g1…g3. Therefore, to avoid any bias, test set instances were excluded from test and training set features list, resulting in a reduced number of features for both test and training sets.

### Class imbalance effect

An undersampling was employed to deal with situations where an imbalance in positive and negative instances was observed. During classifier design, the majority class (negatives) was undersampled in each training fold of five-fold cross-validation. For this purpose, the PercPos method from unbalanced R package was used and perc value was set to 30, resulting in a subset of 30% positive and 70% negative instances. The five-fold cross-validation of classifiers built on undersampled datasets was repeated 20 times and the mean AUC value was reported.

### Constructing functional similarity matrix

Functional similarities between genes were determined using GO annotations. GO is a structured and controlled vocabulary of terms to describe gene products. GO is structured as a Directed Acyclic Graph (DAG) in which terms are represented as nodes and relationships between different terms are represented as edges. The ease of searching and richness in biological information has made GO an imperative resource for studying genes characteristics. Resnik (20), Wang (21), and Relevance (Rel) (22) semantic similarity measures with Maximum (Max) combining strategy were used to construct functional similarity matrices for disease and non-disease genes. Rel and Resnik are Information Content (IC) based measures. Let p(c) be the probability of usage of GO term c in a given GO corpus. The IC of a term can be defined as:

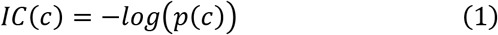

IC-based methods compute a semantic score between two GO terms using the IC of their Most Informative Common Ancestor (MICA) term. Wang measure uses the hierarchical structure of the GO to calculate the semantic similarity scores between the given terms.

The Max strategy of combining semantic similarity scores across all GO terms associated with given genes was used. Let g1 and g2 are two different genes annotated with following list of GO terms (*t*_11_,*t*_12_ ⋯ ⋯ *t*_1*m*_) and (*t*_21_,*t*_22_ ⋯ ⋯ *t*_2*n*_), respectively. The max combining criteria compute the maximum semantic similarity score over all possible pairs of terms between the two term lists as follows:

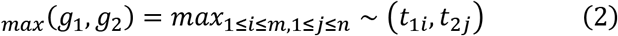

GO terms with electronic evidence code (IEA) were dropped during semantic similarity score calculation. GO information for both ASD and non-mental genes were obtained from org.Hs.eg.db R package (23). Only terms from the biological process aspect were used to assess functional similarities between the genes. All semantic similarity measures were implemented using GOSemSim (version 2.0.4) R package (24). To guarantee that the results are independent of the measure, we decided not to use our in-house developed DiShIn tool and measure (25), but we expect to get similar or better performance with more refined measures.

There are more than 40 thousand GO terms representing biological concepts. However, some genes still lack GO information and result in missing semantic scores. Genes with missing similarity scores were excluded at the beginning of the analysis. We did not assign 0 values to missing scores, to avoid confusion because there were with the genes in dataset that really had a semantic similarity of 0. Therefore, replacing missing scores with 0 can induce bias in predictions and also could result in a bias estimation of classifier performance. All disease gene prediction approaches largely depend upon data and each dataset has its own limitations. For example, protein-protein interaction networks have the limitation of not considering weakly interacting proteins, as these are difficult to ascertain experimentally. Similarly, the presented approach is also dependent upon gene semantic similarities scores, which were unavailable for a few genes. However, in our dataset, semantic scores were missing for only 170 genes (9.5%). Out of which 96 (14.6% of positive instances [disease genes]) belong to the positive class and 74 (6.5% of negative instances [non-disease genes]) were negatives. This small percentage of positive instances with missing similarities will not have a significant effect on classifier performance.

### Automated workflow

To make the designed methodology accessible to researchers lacking extensive machine learning and programming knowledge, we developed an easy to use workflow. The workflow was developed using the KNIME framework (https://www.knime.com/). KNIME is an open source framework implementing several machine learning algorithms and data analysis utilities. The developed automated KNIME workflow incorporates all the above-discussed machine learning and semantic similarity measures to predict disease genes.

### Case study dataset

To implement the proposed methodology, genes with evidence for an involvement in ASD were obtained from the Simons Foundation Autism Research Initiative (SFARI) gene database (N = 990) (https://gene.sfari.org/), accessed on March 2018. The SFARI gene database is a catalogue of ASD relevant genes that have been reported by previous studies. In the SFARI database, genes are scored into seven different categories with respect to the strength of the available evidence. Genes with strongest and reproducible evidence are assigned to category 1 and genes with certain reproducibility limitations are placed in category 2. Category 3 and 4 contains genes with evidence from small studies of ASD candidate genes, while genes belonging to category 5 have indirect evidence of association. Category 6 genes are not associated with ASD. The SFARI gene database also contains syndromic genes that cause genetic pathologies presenting with ASD symptoms. Syndromic genes are placed in a separate syndromic category. Genes from category 1, 2, 3 and 4 (N = 588) were selected for the analysis. All genes classified as syndromic but are not in these categories were excluded. Besides ASD association, syndromic genes are also associated with diverse phenotypes and may disrupt biological processes unrelated to ASD. SFARI genes of category 1 and 2 were designated as High confidence Disease (HD) genes (N = 82), while genes from 3 and 4 SFARI categories were considered as Low confidence Disease (LD) genes (N = 506) (Fig 2). Non-mental genes (N = 1189) were obtained from the Krishnan et al. (10).

**Fig2.**
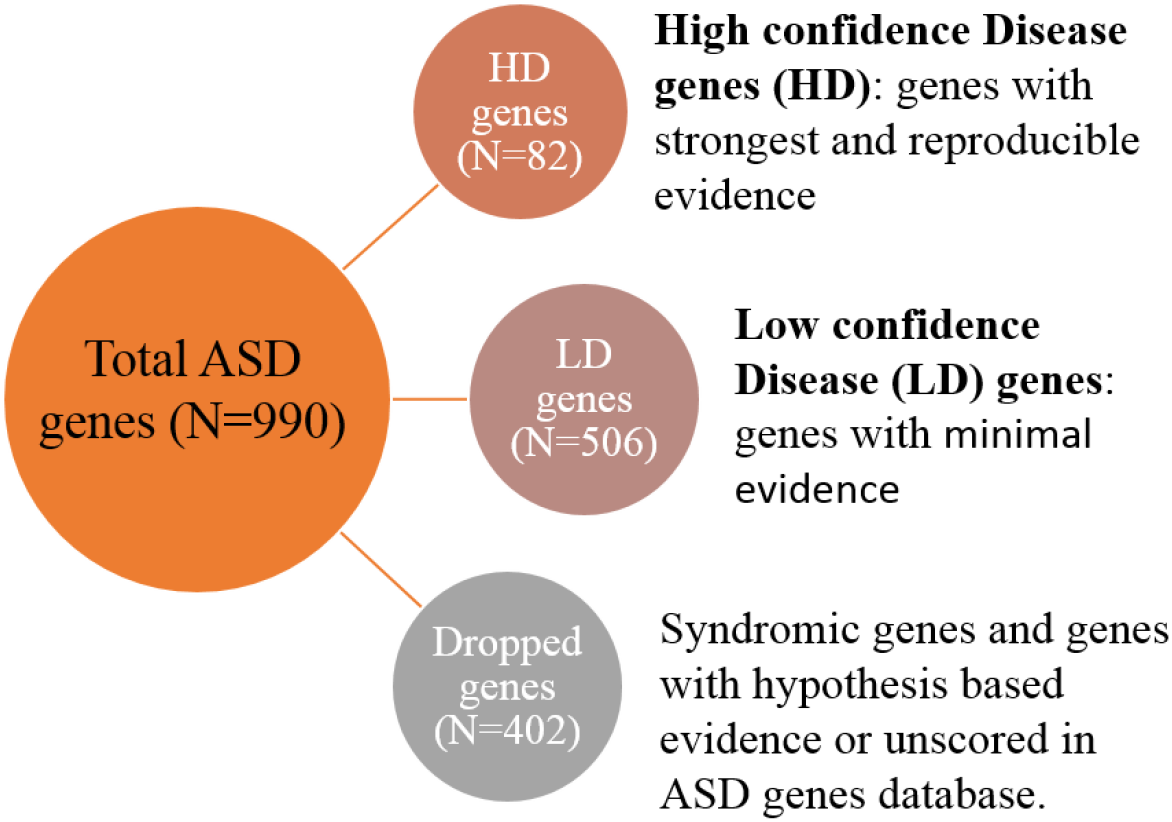
Classifying ASD database genes into HD genes and LD genes using the available evidence strength.

Fig 3 shows the methodology used to predict ASD genes. We modelled the task of predicting disease genes using machine learning methods as a binary classification problem, positive class (+1) for disease genes and negative (-1) for non-disease genes. Therefore, all ASD genes, HD and (HD + LD) composed the positive class, while non-mental genes composed the negative class. Negative class genes were re-examined for their ASD relevance and 49 of them were found in ASD gene database. These genes were considered as ASD genes and were assigned to the positive class.

### Experimental setup

RF, NB, linear-SVM, radial-SVM based classifiers were generated on each semantic similarity matrix (computed using Resnik, Wang, and Rel semantic measures). The classifiers were divided into two different categories depending upon validation strategy and ASD genes classes (HD and LD)

First, classifiers were trained and tested on HD and non-mental genes and were evaluated using stratified five-fold cross-validation. There were less HD ASD genes than non-mental genes. Therefore, to control class imbalance, the majority class (non-mental genes) was undersampled. Secondly, functional similarities of HD +LD and non-mental genes were calculated to design various classifiers. The performance of these classifiers was assessed using two different assessment approaches, namely stratified five-fold cross-validation and held-out restricted five-fold cross-validation.

### Predicting new ASD genes using the semantic similarity based classifier

Copy Number Variants (CNVs) are common genetic risk factors for many diseases, including ASD. Specially, *de novo* CNVs play an important role in ASD pathogenesis and occur at higher rates in patients than in their siblings or controls (26). Mostly, *de novo* CNVs (present in patients but not in their parents) frequently disrupt many genes, posing challenges for the identification of the disease among the many disrupted genes.

ASD candidate gene predictions were made to further estimate the predictability of the built classifier. For this purpose, genes disrupted by rare *de novo* CNVs (N = 80) in ASD patients were obtained from Sander et al. (27). We used our generated classifier to predict ASD genes from genes spanned by rare *de novo* CNVs in subjects that were diagnosed with ASD. Genes overlapping with training set and lacking GO annotations were excluded from analysis.

## Results

In this study, we evaluated the performance of machine learning methods in predicting the ASD candidate genes. For this purpose, functional similarities of ASD genes along with non-mental genes were calculated by applying semantic similarity measures. Table 1 shows the performance of different machine learning classifiers constructed over functional similarities scores, determined by using three different semantic similarity measures.

Resnik, Wang, and Rel measures were used to calculate functional similarities for HD ASD and non-mental genes. Four different machine learning methods (RF, NB, linear-SVM, and radial-SVM) were trained on each semantic similarity matrix, resulting in total of twelve different classifiers for HD genes. Out of these classifiers, RF-based classifiers trained and tested on all semantic similarity matrices outperformed the other classifiers (Table 1). Table 1: The performance of classifiers trained and tested over different semantic similarities matrices and validated using stratified and held-out restricted stratified validation. The performance of the Krishnan et al.’s method in terms of AUC value: unweighted HD genes based classifier = 0.73, weighted HD + LD classifier = 0.74, weighted HD + LD classifier and evaluated with HD genes = 0.80 - 0.89

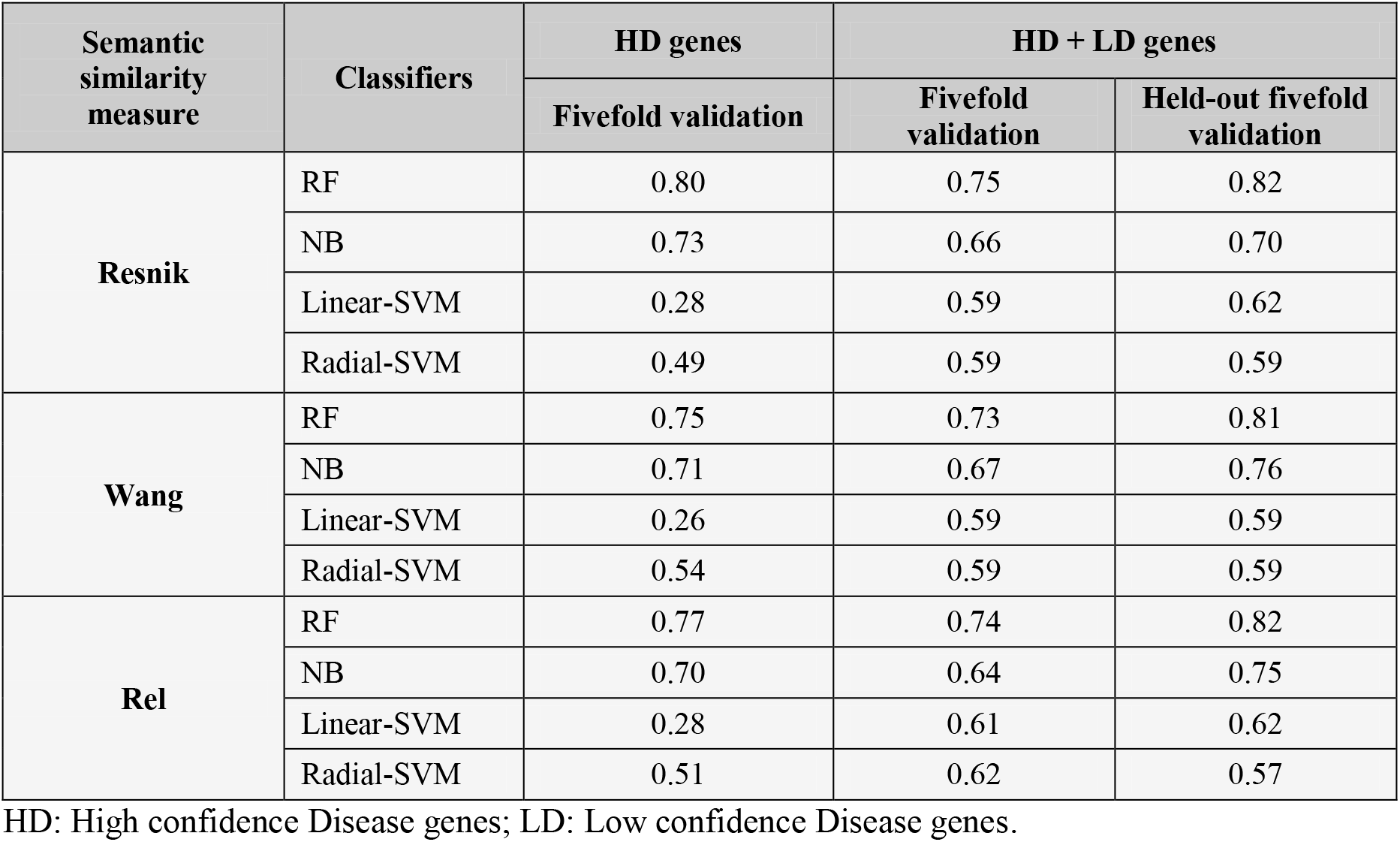

RF classifier trained on Resnik semantic scores obtained best results than the other RF classifiers, although the differences in AUC values were minor, indicating the independence of the methodology to the semantic measure. Linear and radial SVM failed to classify HD ASD genes. The stratified five-fold cross-validation approach was used to evaluate all the classifiers of HD and non-mental genes.

Classifiers were also built using functional similarities of HD + LD and non-mental genes and were evaluated by stratified five-fold cross-validation. The performance of classifiers for each semantic similarity matrix of HD + LD genes and non-mental genes was lower than the HD genes based classifiers, except for linear, and radial SVM (Table 1). RF classifier trained and tested on Resnik semantic scores of HD + LD genes and non-mental genes achieved higher AUC value than the other classifiers of HD + LD genes. Similarly, for HD genes, the performance of RF classifiers trained and tested separately on HD+LD semantic similarities matrices, computed using Wang, Resnik, and Rel measures, was comparable (Table 1).

A held-out restricted validation approach was adapted from Krishnan et al. to assess the performance of the proposed classification approach. For this purpose, classifiers were built using functional similarities of HD + LD genes and non-mental genes. However, during stratified five-fold cross-validation, evaluation was done using only HD ASD and non-mental genes.

For each semantic similarity matrix, held-out restricted RF classifier outperformed HD and HD+LD classifiers (evaluated using stratified five-fold cross-validation). In parallel with previously built classifiers (HD and HD+LD), the performance of held-out restricted classifiers for all semantic similarity measures was comparable (Table 1). Despite the improved performance than HD and HD+LD classifiers, held-out restricted linear and radial SVM failed to classify ASD genes (Table 1). NB classifiers that were evaluated only on HD genes also obtained better performance over other classifiers, evaluated without any restriction. However, improved performance of NB classifiers was only observed for Wang and Rel semantic scores (Table 1).

The reported semantic similar matrices, Resnik, Rel and Wang were calculated using the Max combining criteria. Semantic similar matrices were also calculated by the Boot Mean Average (BMA) combining criteria. However, classifiers constructed using genes semantic scores with the BMA combining criteria showed lower performance than classifiers built on semantic scores of Max combining criteria (S1 Table).

We also have developed a KNIME workflow of the designed methodology. The instantiation of the proposed methodology is described in Fig 4. The workflow comprised four different layers, 1) input layer, 2) functional similarity layer, 3) classifier layer, and 4) output layer. The input layer takes a list of disease and non-disease genes as input. For this list, the functional similarity matrix is calculated in the second layer by selecting a semantic similarity measure (one out of Wang, Resnik, and Rel measures). The classifier layer implements RF, NB, linear-SVM, and radial-SVM machine learning methods with stratified five-fold cross-validation. Lastly, the fourth layer measures the classifier performance by estimating AUC for the generated classifiers. The designed workflow serves as a tool to predict disease genes.

**Fig4.**
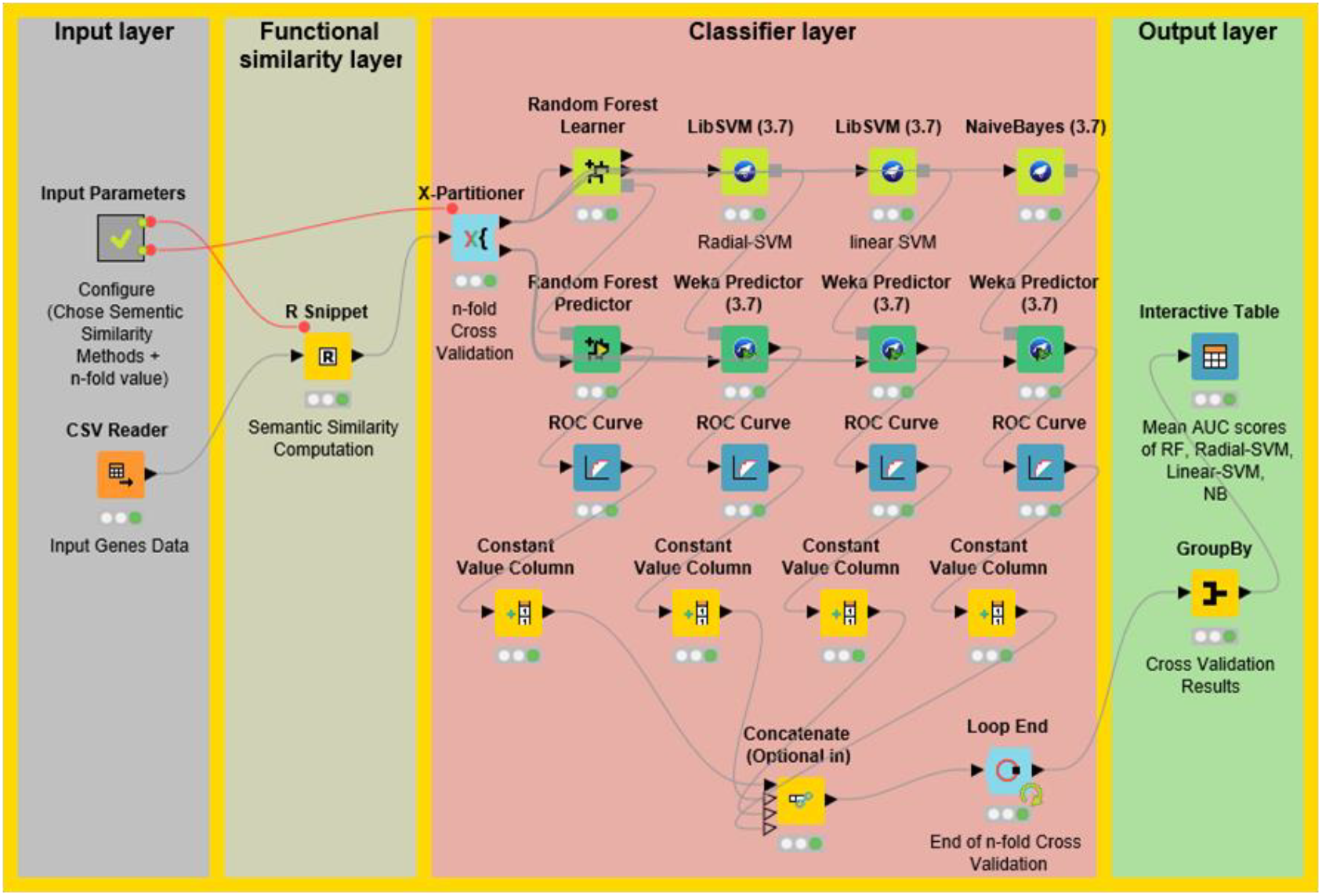
The architecture of the automated workflow for predicting disease genes. Functional similarity layer is the instantiation of methodology step 1, while the classifier layer implements the steps 2 and 3 of proposed methodology (Fig 1).

It can be extended according to the specific objective such as adding another class of disease genes. This workflow also incorporates R nodes to calculate functional similarity between genes, thus allowing programming experts to add any new method by modifying the corresponding R node. Moreover, the model generated using automated workflow can be saved on the local computer and can also be easily invoked to make predictions for new datasets.

To predict novel ASD candidate genes from genes disrupted by de novo CNVs in patients, we used our best classifier, namely the RF classifier trained and tested on Resnik semantic similarity matrix, calculated using HD and non-mental genes. This RF-Resnik classifier was also evaluated by tuning its parameters, such as number of trees. RF classifier showed the highest performance when the number of trees was set to 500 (S2 Table). The classifier predicted 73 ASD genes out of 554 genes that were disrupted by denovo CNVs in ASD patients. The Enrichr (28) tool was employed to find enriched biological pathways and Human Phenotype Ontology (HPO) terms for predicted genes. 99 HPO terms were found significantly enriched for 73 predicted genes (Table 2 and S3 Table).

**Table 2:**
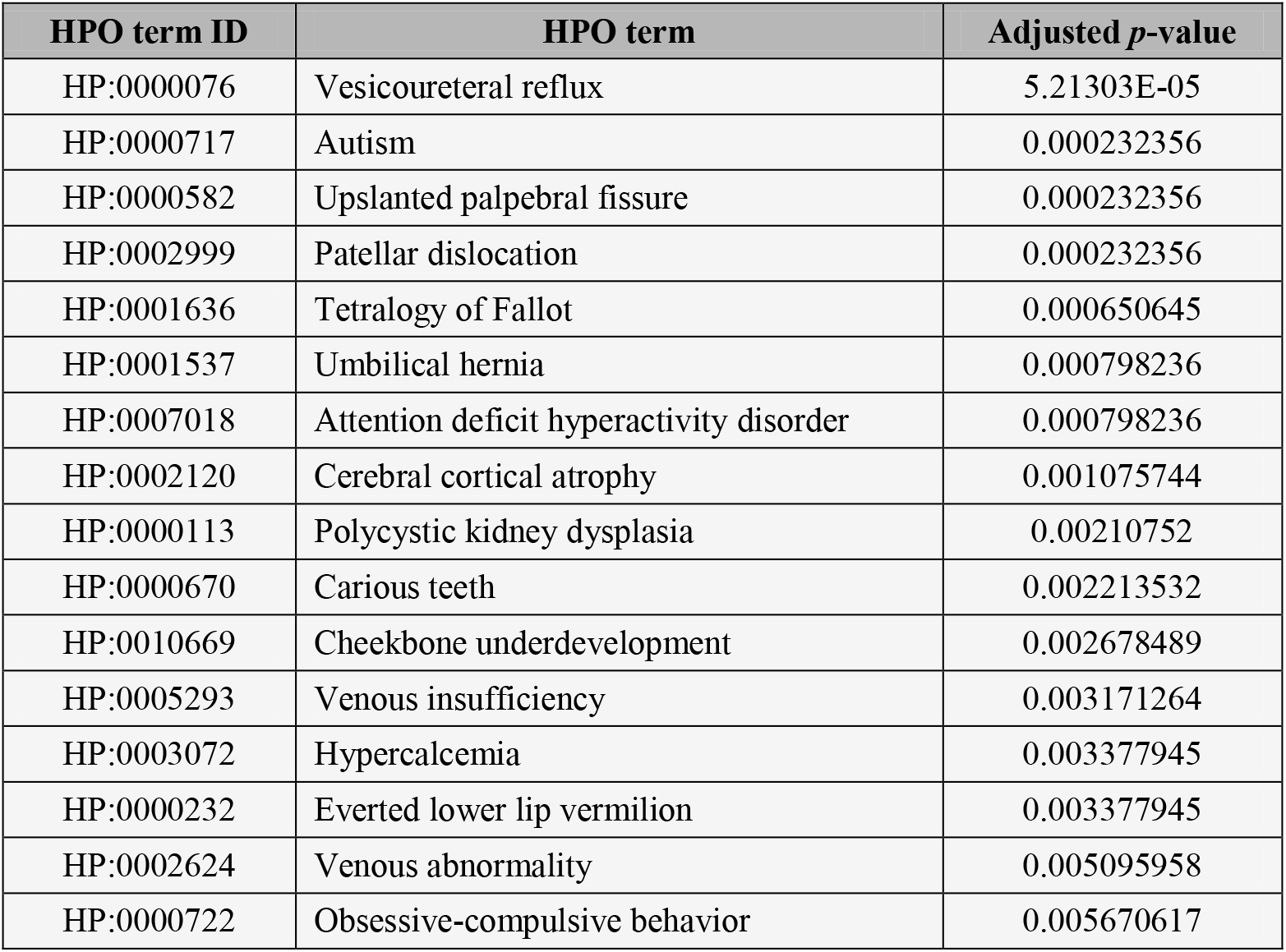
Top 16 enriched HPO terms for predicted ASD genes

The top 2 most significantly enriched HPO terms were *Vesicoureteral reflux* and *Autism*.However, only the histone modifications pathway was marginally significantly enriched for predicted genes (adjusted p = 0.065 at α< 0.05).

## Discussion

Complex diseases exhibit a multifactorial etiology, rendering the identification of biological markers challenging. For example, ASD is a complex neurodevelopmental disorder for which diagnosis relies solely on behavior assessment because efficient biological markers are absent so far. Herein, we reported a machine learning based methodology to predict ASD genes.

Overall, independently from the gene-weighting criteria our RF classifier trained only on gene semantic similarity scores (generated using HD ASD and non-mental genes) obtained higher performance than the Krishnan et al HD ASD genes based classifier, which used protein-protein interaction, gene expression, and gene regulatory information of genes to predict ASD candidate genes (10). Krishnan et al. was selected as state of the art method because to the best of our knowledge was the most recent application of machine learning techniques to identify ASD genes. The study included extensive validation of their methodology and reported real applications in solving a biological problem. We also used the same genes, both ASD and non-ASD genes. Another reason for choosing Krishnan et al. was to compare results from held-out and stratified fivefold cross-validations. The objective was to study which approach is more suitable.

The HD ASD genes based classifier also outperformed the RF classifiers using the semantic similarities of all known ASD (HD + LD) genes. The reduced performance of the classifier trained on HD + LD genes could be explained due to the large number of genes that have weak evidence of involvement in ASD, including, for example, 339 genes from SFARI category 4. Further studies are required to verify the associations of these genes with ASD. Moreover, our classifier trained on HD + LD ASD genes but evaluated on only HD ASD genes as held-out showed better performance than the held-out restricted weighted classifier of Krishnan et al.. However, evaluating classifier performance using only HD ASD genes may provide biased estimates. ASD presents a wide spectrum of clinical phenotype. The classifier tested on only HD ASD genes may lack robustness for genes that are responsible for a milder expression of ASD or for comorbidities, thus limiting the coverage of the predictive classifier.

The strength of the reported semantic similarity scores is its independence from the features (genes) weighting scheme, defined by the ranking of genes according to available evidence. The standard criteria for ranking ASD genes by available reference is not yet completely defined (29), and, therefore, it is possible that incorporating weighting criteria for genes could introduce a bias. In the present study, we used semantic similarity scores of genes, which range from 0 to 1, with semantic similarity scores closer to 1 indicating high functional similarity between genes. If the gene is not directly involved in ASD then the target gene will show less similarity with disease-causing genes, hence lowering its contribution in classification. For this reason, we did not apply any weighting criteria. This automates the process and allows increasing performance as GO evolves.

The improved performance of the classifier with semantic similarities further confirmed the importance of GO annotations in predicting disease genes. GO provides a consistent and deeper level of biological information than the protein-protein interactions or genes expression profiles. Functional information from GO structure can overcome the shortcomings of protein-protein interactions and expression profiles because GO annotations are less dependent on context. Additionally, our study further confirmed that machine learning methods can be used to predict disease genes and their performance can be improved by adding more data with different domain information.

One limitation of this approach is the full dependence on the available annotation resources. Genes lacking GO annotations are filtered in the analysis. But this may improve in future because the number of GO terms and annotations are continuously increasing with time.

We also provided a downloadable and easy to use workflow of the designed methodology. The designed workflow permits the user to test more than one machine learning and functional similarity methods. The workflow allows domain-specific experts to replicate the reported methodology to identify disease-causing genes and use their knowledge to further improve the predictions by tuning the parameters according to their need. The developed workflow includes all necessary tools for disease gene prediction and provides an opportunity for researchers to analyze their own private data without uploading it to external websites, thus preventing any privacy issues.

Enrichment analysis revealed the enrichment of ASD predicted genes in *histone modification* pathway. This was consistent with previously published studies, reporting histone genes involvement in ASD (30). Only one pathway was significantly enriched for ASD predicted genes. This might be because of a small number of input ASD genes. Prediction with larger number of genes might produce more statistically significant pathways.

ASD core symptoms are defined by deficits in social interaction and communication, and repetitive behavior. However, these core symptoms manifest with a wide range of phenotype, indicating ASD phenotypic heterogeneity. Functional enrichment analysis of ASD genes, predicted by the RF classifier also showed a wide range of phenotypes. ASD predicted genes were enriched in *Autism* and *obsessive-compulsive behavior* phenotypes, further confirming the feasibility of the classifier in predicting new ASD genes. In addition, five genes were also associated with the *attention deficit hyperactivity disorder* HPO term. Attention deficit hyperactivity is one of most the common ASD comorbidity. A large percentage of ASD children also develop attention deficit hyperactivity disorder symptoms (31).

Although Random Forest (RF) performed better than other methods, its performance was comparable with Naive Bayes (NB) algorithm. Therefore, we do not expect that RF will achieve the best performance for other complex disease genes as well. The Support Vector Machines (SVMs), both linear and nonlinear failed to make reasonable predictions, probably, they are sensitive to number of features and number of records. Also, SVMs perform preprocessing of data, which may not be feasible for this dataset.

In this study, we reported that gene functional similarities, assessed using GO annotations can be used to identify new ASD genes. The classifier trained and tested using HD ASD genes outperformed all others and previously reported classifiers. The objective of this work was not to find the best semantic similarity measure for calculating functional similarities of genes. However, the performance of the classifier can be further improved by including more refined semantic similarity methods. Additionally, text mining techniques may also be employed to find the functionality of genes in literature that are lacking semantic scores (32). Moreover, to attain reliable ASD genes predictions, future studies should be focused on combining protein-protein interactions with semantic similarity scores. Additionally, integration of information such as gene expression and pathways data with semantic scores can further improve the power of prediction.

In summary, we validated our hypothesis and have shown that the performance of available state of the art methods for complex disease gene prediction can be improved by using only GO annotations. One novel aspect of presented manuscript was the comparison of stratified and held-out restricted cross-validation. The results indicated that classifiers built on GO annotations did not require held-out restricted classifiers. Approximately equivalent performance to ASD held-out restricted classifiers can be achieved by training machine learning methods, validated using five-fold cross-validation, on semantic similarity scores of only high confidence ASD disease genes. This work demonstrated that machine learning methods are a feasible approach to dissect the genetic heterogeneity of complex diseases such as ASD and to find novel disease genes. The results of this study can also further assist in designing genetic screening and lab experiments on ASD genetic risk factors.

## Acknowledgements

We would like to thank Ana Rita Marques and André Lamúrias for their valuable suggestions to improve this work. Funding: DeST: Deep Semantic Tagger FCT funded project PTDC/CCI-BIO/28685/2017.

